# The significance of *sukumo* (composted indigo leaves) as a microbial source for traditional Japanese indigo dyeing

**DOI:** 10.64898/2026.01.29.702489

**Authors:** Souichiro Kato, Kensuke Igarashi, Shusei Kanie, Kyosuke Yamamoto, Wataru Kitagawa, Takashi Narihiro, Kenta Watanabe, Kenta Fujii, Isao Yumoto, Yoshiyuki Ueno

## Abstract

A critical process in traditional Japanese indigo dyeing is the microbial reduction of indigo within the dye suspension, which consists solely of *sukumo* (fermented indigo leaves), wood ash lye, and microbial nutrients such as wheat bran. Although *sukumo* has long been recognized as a potential microbial source, the microbial community dynamics during its production is still largely unexplored, and its contribution to dyeing performance via microbial supply remains poorly characterized. In this study, we investigated the significance of *sukumo* as a microbial source, in addition to its established role as a pigment source. We conducted a time-series analysis of microbial communities throughout the four-month fermentation process of *sukumo*, using weekly samples collected from two geographically distinct indigo dyeing studios. Microbial profiling revealed similar successional patterns between the two sites. Notably, in the later stages of fermentation, known indigo-reducing bacteria, such as the genus *Oceanobacillus* and the family *Tissierellaceae*, emerged at both locations. Laboratory-scale dyeing experiments using immature *sukumo* demonstrated that supplementation with a small amount of mature *sukumo* restored dyeing activity and increased the abundance of *Oceanobacillus* and *Tissierellaceae*. Furthermore, the addition of the indigo-reducing isolate *Tissierellaceae* strain TU-1 to the immature *sukumo*-based dye suspension led to a marked enhancement in dyeing performance. These findings highlight the critical role of *sukumo* as a microbial source in traditional indigo dyeing and suggest that prolonged fermentation is essential for nurturing functional indigo-reducing bacteria. This insight provides a foundation for improving dye suspension performance through targeted microbial community management.

## Introduction

As the name ‘Japan Blue’ suggests, indigo dyeing is a traditional technique that has been widely practiced in Japan for textile coloration (Ueno, 2023). Similar to bioprocesses used in the production of fermented foods and beverages, traditional indigo dyeing effectively harnesses the functions of specific microorganisms (Aino *et al*., 2010). The process begins with the manufacturing of a dye material known as *sukumo* (Fig. 1), which is produced by composting the leaves of *Polygonum tinctorium*, a plant rich in indican (indoxyl-β-D-glucoside, a precursor of indigo), for a period of up to four months. To produce the indigo dye suspension, *sukumo* is suspended in water alkalized with wood ash and slaked lime, and supplemented with microbial nutrients such as wheat bran. The mixture is then fermented for one to two weeks, during which microbial activity renders the solution suitable for dyeing. In the dye suspension, microorganisms reduce indigo, which is insoluble in alkaline water, to its soluble form, leuco-indigo. Textiles are dyed by immersing them in the dye suspension containing leuco-indigo, which is subsequently oxidized by air to regenerate indigo on the fabric. Numerous studies have investigated the microorganisms present in indigo dye suspensions and the dynamics of microbiota during the fermentation process, using both culture-dependent and -independent methods (Nishita *et al*., 2017; Aino *et al*., 2018; Tu *et al*., 2019; Lopes *et al*., 2021; Tu *et al*., 2021; Li *et al*., 2022; Farjana *et al*., 2023). In particular, many species of indigo-reducing bacteria have been isolated, and their physiological characteristics have been examined (Nicholson and John, 2005; Park *et al*., 2012; Hirota *et al*., 2013a; 2013b; Nakagawa *et al*., 2022; 2024; Tu and Yumoto, 2025). However, despite the fact that microbial activity is also essential in the production of *sukumo*, no microbiological studies of the *sukumo* production process have been reported to date.

**Fig. 1.**
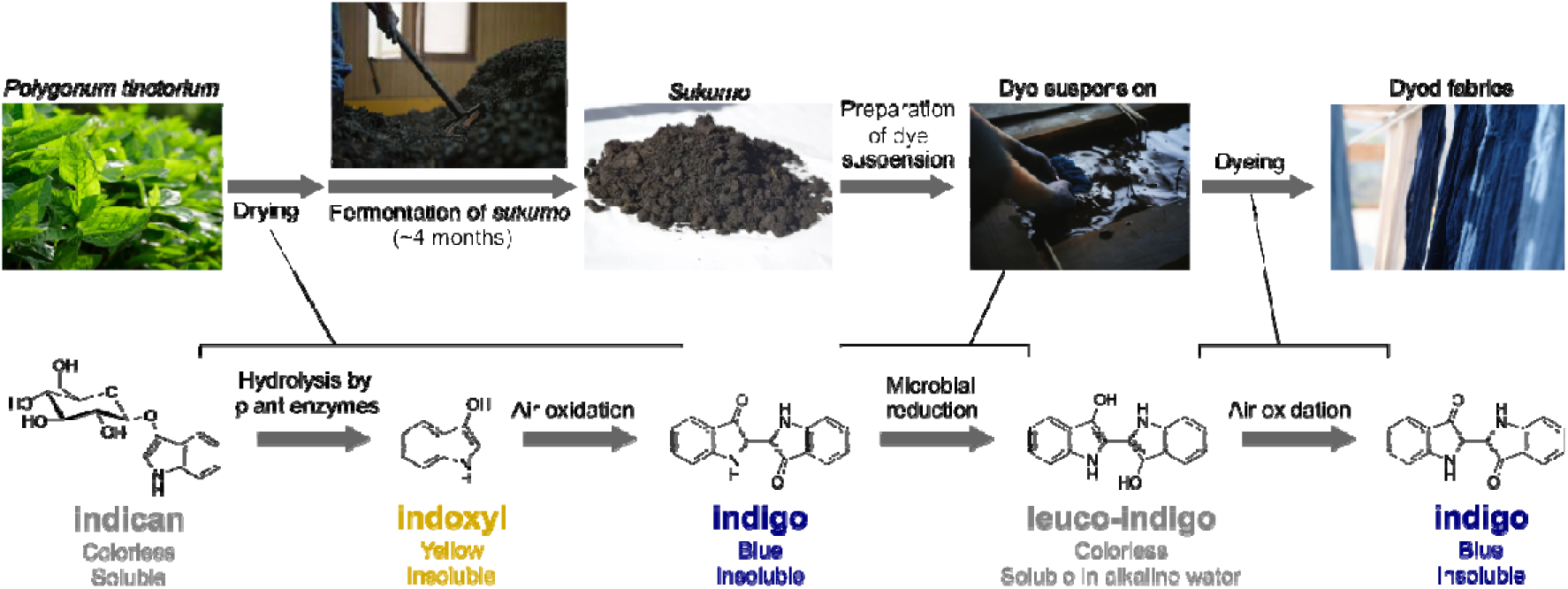
Schematic representation of the traditional indigo dyeing process and the associated chemical transformations of indigo and its derivatives. The color and water solubility of each compound are indicated.

The production of *sukumo* begins with the cultivation of indigo plants. After harvesting, the plants are sun-dried, cut, and only the leaf portions are collected. The dried leaves are then piled and moistened to initiate microbial fermentation. Throughout the *sukumo* production process, the piles are rehydrated and thoroughly mixed once a week. Similar to general composting processes, microbial activity causes the temperature within the piles to rise, sometimes reaching up to 80 °C. As fermentation progresses, an ammonia odor is released, likely due to the degradation of proteins, and the pH gradually increases. After three to four months of fermentation, the pH reaches 9 to 10, marking the completion of *sukumo* production. The *sukumo* production process plays a critical role in decomposing plant-derived organic matter, thereby reducing its volume and improving both portability and preservation. In addition, because *sukumo* serves as a primary microbial source for dye suspensions, it is expected to provide beneficial microorganisms, including indigo-reducing bacteria. However, to date, no studies have investigated microbial dynamics during *sukumo* production, and the significance of *sukumo* as a microbial source has yet to be demonstrated.

In this study, we analyzed microbial dynamics during *sukumo* production over a four-month fermentation period in two geographically distinct regions. Furthermore, the role of *sukumo* as a microbial source was evaluated through dye suspension culture experiments using incompletely fermented, immature *sukumo*, along with assessments of the effects of externally introduced microbial sources, specifically, commercially available mature *sukumo* and a pure culture of an indigo-reducing bacterium.

## Materials and Methods

### Bacterial strains and culture conditions

*Tissierellaceae* strain TU-1 (JCM 37169) was cultured anaerobically in PYA (peptone/yeast extract/alkaline) medium at 30 °C without shaking. The PYA medium was prepared as follows: vials (68 mL capacity) were filled with 18 mL of PY solution (8.9 g peptone, 3.3 g yeast extract, and 1.1 g K_2_HPO_4_ per liter), flushed with N_2_ gas, and sealed with butyl rubber stoppers and aluminum caps prior to autoclaving. After autoclaving, 2 mL of filter-sterilized Na_2_CO_3_/NaHCO_3_ buffer (1 M, pH 10) was added.

### Production processes of sukumo and sampling procedures

This study used *sukumo* produced at two geographically distinct indigo dyeing studios in Japan. One was produced at the studio of Aiya-terroir (Fukuyama, Hiroshima, 34.6569°N, 133.3823°E) (hereafter referred to as T *sukumo*), and the other at the studio of Watanabe’s Co. Ltd. (Kamiita, Tokushima, 34.1046°N, 134.4047°E) (hereafter referred to as W *sukumo*). Both studios use leaves of *Polygonum tinctorium* as raw material. The dried leaves are piled and moistened with water, then thoroughly mixed once per week throughout the fermentation process. Fermentation of W *sukumo* began with 720 kg of dried leaves, with an additional 180 kg added during each of the first, second, and third weeks of mixing, resulting in a total of 1,260 kg. T *sukumo* was produced using a total of 1,110 kg of dried leaves. Fermentation began with 450 kg, followed by the addition of 120 kg in each of weeks 1 through 4, and 90 kg in each of weeks 5 and 6. The *sukumo* production processes were completed within 16 weeks. Samples for analysis were collected weekly after each mixing operation, in duplicate at each time point. Samples for culture experiments were immediately transported to the laboratory and stored at 4 °C until use. Samples for chemical and microbial community analyses were stored at −20 °C until use.

### *Quantification of indigo in* sukumo

Chemically reducible indigo in *sukumo* was quantified as an estimate of the indigo available to microorganisms. Approximately 20 mg (dry weight) of *sukumo* was suspended in 5 mL of hydrosulfite solution (17.4 g L^-1^ Na_2_S_2_O_4_ in 1 g L^-1^ NaOH), mixed by vortexing, and incubated at 75 °C for 10 min to reduce indigo to leuco-indigo. The suspension was then centrifuged at 12,000 × *g* for 3 min, and the supernatant containing leuco-indigo was collected. To oxidize leuco-indigo to water-insoluble indigo, the supernatant was acidified by mixing it with an equal volume of 2 M HCl. The mixture was centrifuged again at 12,000 × *g* for 3 min, and the resulting precipitate was dissolved in dimethyl sulfoxide (DMSO). The concentration of indigo in the DMSO solution was determined using ultra-performance liquid chromatography (UPLC; ACQUITY UPLC H-Class system; Waters, MA, USA) equipped with an ACQUITY UPLC BEH C18 column ( 2.1 × 100 mm, 1.7 μm; Waters) and a multiwavelength detector (ACQUITY UPLC PDA eλ detector; Waters). The HPLC conditions were as follows: mobile phase, aqueous acetonitrile containing 0.1 % (v/v) formic acid; gradient conditions, 10–30 % aqueous acetonitrile containing 0.1 % (v/v) formic acid over 10 min, 30–100 % over 5 min, 100 % for 5 min; flow rate 0.2 mL min^−1^; UV detection, 612 nm; column temperature, 30 °C; injection volume, 5 μL. Standard curves were generated using synthetic indigo powder (Sigma-Aldrich, Burlington, MA, USA) dissolved in DMSO.

### Microbial community analysis

DNA was extracted using FAST DNA Spin Kit for Soil (MP Biomedicals, Irvine, US) and then purified using QIAquick PCR Purification Kit (Qiagen, Hilden, Germany). PCR amplification of V4 region of bacterial and archaeal 16S rRNA gene was conducted using the purified DNA as a template, KAPA HiFi HotStart ReadyMix (Roche Sequencing Solutions, Pleasanton, CA, US), and primers 515’F/806R (Hugerth *et al*., 2014). The thermal conditions were as follows: initial thermal denaturation at 95°C for 3 min, followed by 25 cycles of heat denaturation at 95°C for 30 sec, annealing at 55°C for 30 sec and extension at 72°C for 30 sec; and a final cycle at 72°C for 5 min. The PCR product was purified by agarose gel electrophoresis followed by gel extraction using QIAquick PCR purification kit (Qiagen).

The purified PCR product was used as the template for the subsequent index PCR reaction to incorporate a dual-index Nextera barcode and the remaining Illumina adapter sequence (Illumina, San Diego, CA, USA). The PCR condition was the same as above, but the amplification cycle was limited to 8 cycles. The resulting index PCR product was subjected to agarose gel electrophoresis to confirm length of amplified fragments and purified as above. Each PCR product was quantified using the Qubit 4 and its dsDNA BR Assay Kit (Thermo Fisher Scientific, Waltham, MA, USA) and then mixed to prepare amplicon DNA library. The library was subjected to sequencing analysis with an iSeq 100 sequencing system according to the manufacturer’s instructions, utilizing an Illumina iSeq™ 100 i1 Reagent v2 kit (Illumina). The bioinformatics analysis was performed using the QIIME 2 pipeline v.2024.10 (Bolyen *et al*., 2019). The raw sequence data in the FASTQ format were imported into QIIME 2 and demultiplexed. Quality filtering was performed using the q2-demux plugin, and sequence denoising was carried out using DADA2. The taxonomic classification of the amplicon sequence variants (ASVs) was performed using the classify-consensus-blast classifier of the q2-feature-classifier plugin against the SILVA v.138.99 database for 16S ASVs (https://docs.qiime2.org/2024.10/data-resources/). Principal component analysis was performed using Principal Component Analysis Calculator available at Statistics Kingdom (https://www.statskingdom.com/pca-calculator.html).

### *Preparation of indigo dye suspension cultures using immature* sukumo

Two grams of immature *sukumo* were suspended in 49 mL of the lye solution in a 50 mL tube. The lye solution was prepared by suspending 20 g of diatomite powder in 1 L of distilled water, boiling the mixture for 5 min, allowing it to cool to room temperature, and collecting the supernatant. The pH was adjusted to 10 by adding Ca(OH)_2_, and pH adjustments were performed every few days during incubation. Wheat bran (20 mg) was added as a microbial nutrient at weeks 0, 2, and 4 of the culture periods. Cultures were incubated at 30 °C without shaking. For experiments involving the addition of an external microbial source, 0.1 g of commercially available *sukumo* (high-quality *sukumo* indigo, Sato Awa-Ai Factory, Tokushima, Japan) was added to the dye suspension cultures. For experiments involving an indigo-reducing bacterium, 1 mL of culture solution of *Tissierellaceae* strain TU-1 was added. For the control dye suspension cultures, 1 mL of sterilized PYA medium was added. Samples for microbial community analysis were collected weekly from week 1 to week 6. To assess dyeing performance, a small piece of cotton fabric was immersed in the dye suspension for 20 sec, exposed to air, rinsed three times with water, and air-dried. Dyeing intensity was quantified by comparing the grayscale color intensity of dyed and undyed fabrics using photographic data analyzed with the Color Picker tool (https://colorpicker.tools/). The weighted average method was used to convert from the RGB color codes to the grayscale values (0.2989×R + 0.5870×G + 0.1140×B).

## Results and Discussion

### *Production of* sukumo *in two indigo-dyeing studios*

In this study, we used two types of *sukumo* (W *sukumo* and T *sukumo*) produced under different geographical conditions and production methods (see Materials and Methods). The most notable difference lies in the method of adding the raw material, dried indigo leaves. Compared to W *sukumo*, T *sukumo* had a longer period to add new dried leaves, and the relative amounts were larger. The pH transitions during the *sukumo* fermentation processes (Fig. 2A) showed that the pH increased more slowly in T *sukumo*. This observation suggests that the progression of fermentation was slower in T *sukumo*, likely due to the differences in the manufacturing processes described above.

**Fig. 2.**
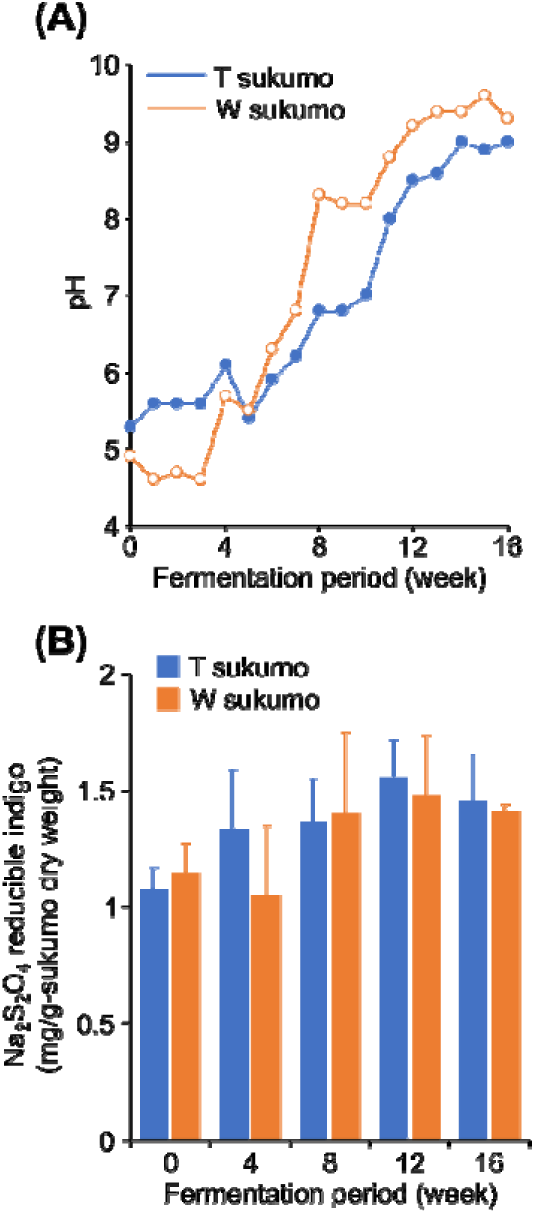
Chemical properties of sukumo during the production process. (A) Temporal changes in pH. (B) Temporal changes in the amount of indigo extractable by reduction with Na_2_S_2_O_4_. Data are presented as the means of three independent samples. Error bars represent standard deviations.

The amount of indigo that can be solubilized by reduction with hydrosulfite (Na_2_S_2_O_4_) was determined as an indicator of the amount of indigo available for microbial reduction (Fig. 2B). The amount of chemically reducible indigo per dry weight of *sukumo* showed an increasing trend during the production process, although the increase was modest, ranging from approximately 1.2 to 1.4 times. It has generally been considered that reducing the amount of plant-derived organic matter while increasing the relative indigo content is one of the key aspects of *sukumo* production. However, the results suggest that this effect is not as pronounced as previously thought.

### *Dynamics of microbiota during the production of* sukumo

Microbial community analysis based on amplicon sequencing of 16S rRNA genes was performed to investigate the variation in microbiota during the *sukumo* production process. The changes in the relative abundances of the dominant phylotypes are shown in Fig. 3A. The PCA based on the microbial community structures indicated that the microbiota in the two different *sukumo* production processes were very similar at the beginning and end of fermentation, with slight differences observed during the intermediate stages (Fig. 3B). Given that they produced *sukumo* in distant locations and by different procedures, it was unexpected that the final microbiota would be so similar.

**Fig. 3.**
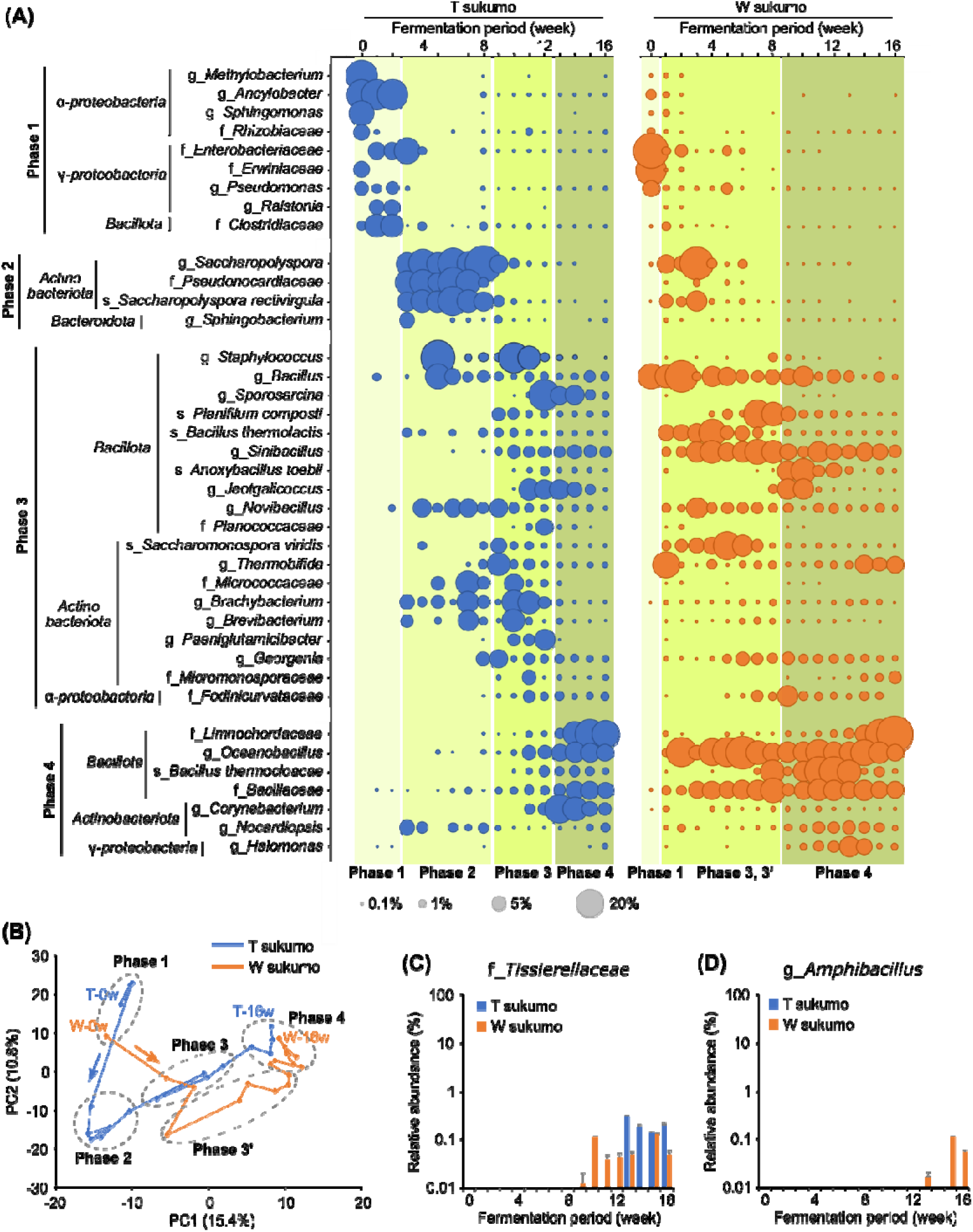
Microbial community analysis during the *sukumo* production process. (A) Temporal changes in the relative abundance of dominant phylotypes (defined as those comprising >5% in at least one sample). (B) Principal component analysis based on the microbial community profile data. ‘T-0w’ represents ‘0-week sample of T *sukumo*’. (C, D) Temporal changes in the relative abundance of f_*Tissierellaceae* and g_*Amphibacillus*. All data represent the average values from two independent samples.

Based on the PCA result and the pH profile, we classified the *sukumo* production processes into four phases (Fig. 3B). Phase 1 represents the early stage of fermentation, encompassing weeks 0 to 2 for T *sukumo* and week 0 for W *sukumo*. *Sukumo* in Phase 1 was predominated by *Alphaproteobacteria*, such as the genera *Methylobacterium* and *Ancylobacter*, and *Gammaproteobacteria,* such as the families *Enterobacteriaceae* and *Erwiniaceae*. Since these bacteria are commonly detected on plant surfaces (de Maayer *et al*., 2014; Iguchi *et al*., 2015; Janda and Abbott, 2021; Agafonova *et al*., 2023), the *sukumo* microbiota in Phase 1 are considered to be strongly influenced by those of dried leaves. Phase 2 is observed only in T *sukumo* (weeks 3 to 8), likely reflecting the slow progress of fermentation due to the frequent addition of dried leaves. Phase 2 is characterized by the dominance of *Actinobacteriota*, particularly three phylotypes belonging to the family *Pseudonocardiaceae* (classified as g_*Saccharopolyspora*, f_*Pseudonocardiaceae*, and s_*Saccharopolyspora rectivirgula*, under the SILVA taxonomy), which are commonly detected from composting processes (Hayakawa *et al*., 2010; Riahi *et al*., 2022). Phase 3 consists of weeks 9 to 12 for T *sukumo* and weeks 1 to 8 for W *sukumo*, with the pH reaching 8 in the final weeks. Bacteria in the phylum *Bacillota*, particularly the families *Bacillaceae* (g_*Bacillus*, g_*Sinibacillus*, and g_*Anoxybacillus*), *Staphylococcaceae* (g_*Staphylococcus* and g_*Jeotgalicoccus*), *Planococcaceae* (g_*Sporosarcina*), and *Thermoactinomycetaceae* (g_*Planifilum*), were dominant in Phase 3, which are also common microorganisms found in composting processes (Haruta *et al*., 2005; Watanabe *et al*., 2008; Han *et al*., 2013; Yang and Zhou, 2014; Wei *et al*., 2018; Liu *et al*., 2022; Dobrzyński *et al*., 2023).

Phase 4 represents the final stage of fermentation, characterized by the pH exceeding 8 and gradually increasing. This phase corresponds to weeks 13 to 16 for T *sukumo* and weeks 9 to 16 for W *sukumo*. In Phase 4, phylotypes belonging to the phylum *Bacillota*, different from those in Phase 3, were dominant. The phylotype belonging to the family *Limnochordaceae*, moderately thermophilic bacteria commonly detected in composts (Watanabe *et al*., 2015; Zhou *et al*., 2023), was the most dominant in the final phase. *Limnochordaceae* was also identified as the predominant bacteria in commercial *sukumo*, but its abundance was reduced in the indigo dye suspension (Tu *et al*., 2021). Several phylotypes belonging to the family *Bacillaceae* (f_*Bacillaceae*, s_*Bacillus thermocloacae*), which are often detected in composts (Watanabe *et al*., 2008), were also predominant in this phase. These *Bacillota* bacteria are expected to contribute to composting in the alkaline environments of *sukumo* production. As expected, phylotypes predicted to be indigo-reducing bacteria were also dominant. The genus *Oceanobacillus* is commonly detected in *sukumo* and indigo dye suspensions (Lopes *et al*., 2021; Tu *et al*., 2021; Farjana *et al*., 2023), and isolates with indigo-reducing ability have been obtained (Hirota *et al*., 2013b; 2013d). Although the abundances of the phylum *Actinomycetes* were generally reduced in Phase 4, a phylotype within the genus *Corynebacterium* became dominant only in T *sukumo*. The genus *Corynebacterium* has often been detected in indigo dyeing suspensions (Aino *et al*., 2010; Tu *et al*., 2021), and a strain with high indigo reduction capacity has been isolated (Nakagawa *et al*., 2022). In addition, several phylotypes closely related to known indigo-reducing bacteria were detected exclusively in Phase 4, albeit as minor populations. The phylotype belonging to the family *Tissierellaceae* was detected in Phase 4 of both T and W *sukumo*, with a maximum abundance of only 0.3% (Fig. 3C). The *Tissierellaceae* bacteria has been consistently identified as a dominant species in nearly all indigo dyeing suspensions (Aino *et al*., 2010; Lopes *et al*., 2021; Tu *et al*., 2021; Farjana *et al*., 2023). Its isolated strains exhibit higher indigo-reducing activity compared to other known indigo-reducers, suggesting that *Tissierellaceae* bacteria plays a critical role in the indigo dyeing process (Tu and Yumoto, 2025). The phylotype within the genus *Amphibacillus*, also known as indigo-reducing bacteria (Hirota *et al*., 2013a; 2013c), was detected exclusively in Phase 4 of W *sukumo*, with a maximum abundance of 0.1% (Fig. 3D). It has been reported that even when indigo-reducing bacteria become dominant in dye suspension, they may be undetectable or present only in trace abundances in the microbial source, *sukumo* (Tu *et al*., 2021). As described below, this study also showed that microorganisms not detected in *sukumo* (*i.e.*, abundances of <0.01%) could become dominant in dye suspensions. The gradual increase in indigo-reducing bacteria toward the final stages of the production process is likely one of the key reasons why *sukumo* requires long-term fermentation, even though indigo reducing bacteria constitute only a minor population. The prolonged fermentation likely facilitates the decomposition of organic matter and a gradual increase in pH, creating an environment favorable for the proliferation of anaerobic alkaliphiles, including indigo-reducing bacteria.

### *Effects of an additional microbial source (commercial* sukumo*) on dye suspensions prepared with immature* sukumo

We conducted laboratory culture experiments to test the following two hypotheses: (1) the staining activity of dye suspensions prepared with immature *sukumo* may be limited due to the absence of key microorganisms, and (2) supplementation with additional microbial sources may improve their staining performance. Dye suspensions in small scales (50 mL) were prepared using immature *sukumo* collected during the final week of Phase 3 of the fermentation process, specifically, week 12 of T *sukumo* (T-12w) and week 8 of W *sukumo* (W-8w). To evaluate the effect of additional microorganisms essential for active dyeing, the cultures further supplemented with commercially available mature *sukumo* (at 1/20 the volume of the immature *sukumo*), referred to as the +*sukumo* series, were also prepared.

The results of staining activity measurements and microbial community analyses of dye suspensions prepared with immature *sukumo* are presented in Fig. 4. Although cotton fabrics were stained in dye suspensions using only immature *sukumo*, more stable staining was achieved in those further supplemented with commercial *sukumo* (Figs. 4A and 4D). Figures 4B and 4E show the quantified color intensities of the stained fabrics. The addition of commercial *sukumo* resulted in a significant increase in staining intensity at weeks 4, 5, and 6 in cultures using T-12w, and at weeks 1, 2, and 6 in those using W-8w. Given that the amount of commercial *sukumo* added was only 1/20 the volume of the immature *sukumo*, the observed enhancement in staining intensity is likely attributable to changes in the microbial communities introduced by microorganisms from the commercial *sukumo*.

**Fig. 4.**
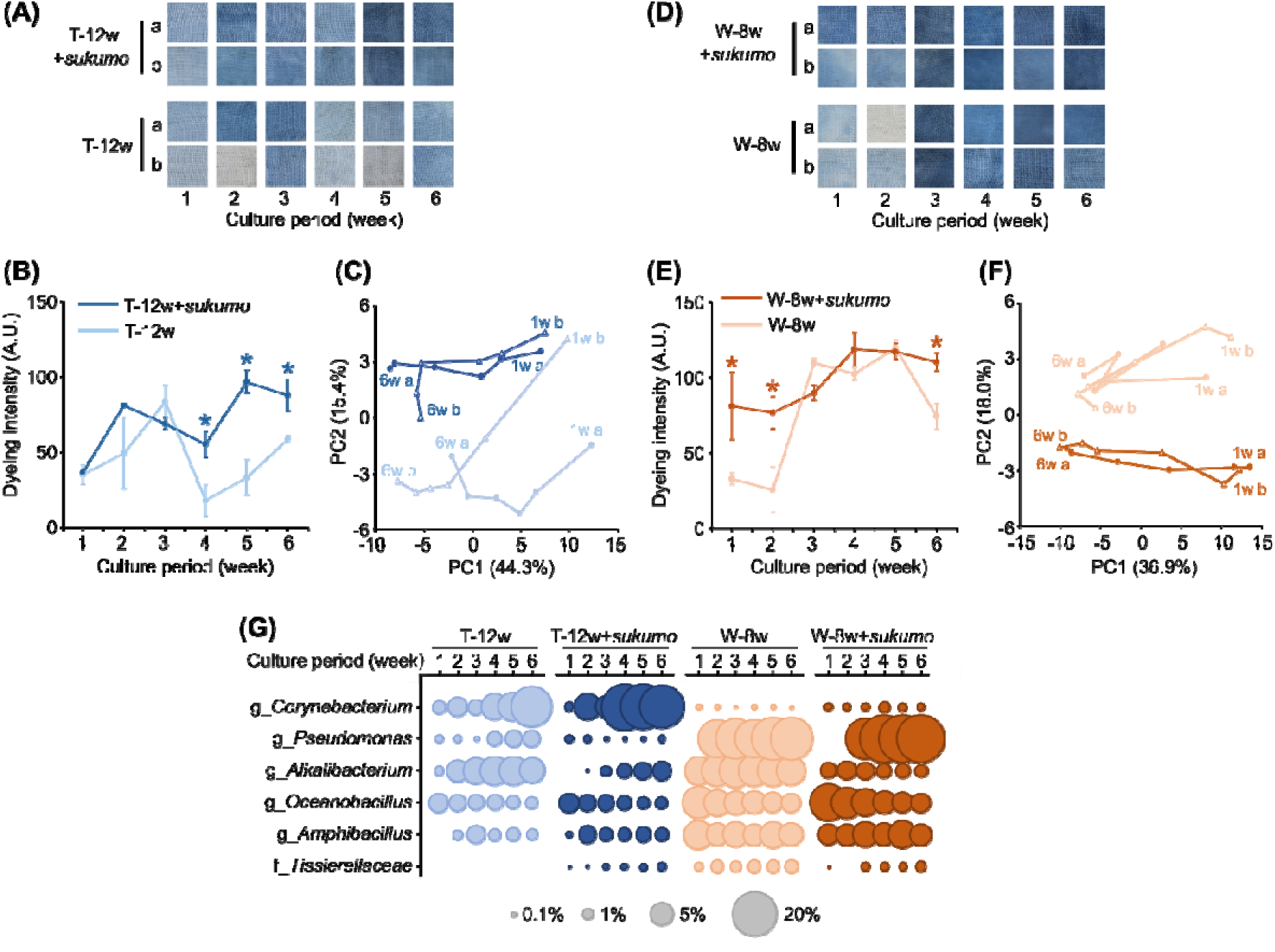
Culture experiments of dye suspension using immature *sukumo* to evaluate the effect of supplementation of commercial *sukumo* as a microbial source. Immature *sukumo* samples were collected from the production process immediately after the pH reached 8. T-12w (A–C) and W-8w (D–F) refer to dye suspension cultures using 12-week of T *sukumo* and 8-week of W *sukumo*, respectively. T-12w+*sukumo* and W-8w+*sukumo* indicate cultures supplemented with a 1/20 volume of commercially available mature *sukumo* as a microbial source. (A, D) Photographs of cotton fabrics stained using each dye suspension cultures. Labels ‘a’ and ‘b’ represent two independent culture replicates. (B, E) Quantification of color intensity based on the dyed fabrics shown in (A) and (D). Asterisks indicate significantly higher color intensity in the +*sukumo* cultures, which was assessed by Student’s *t*-test. (C, F) Principal component analysis based on microbial community profiles of the dye suspension cultures, which are presented in Supplementary Fig. S1. Data points corresponding to weeks 1 and 6 of each culture are annotated in the figure. (G) Temporal changes in the relative abundance of dominant phylotypes (> 1% relative abundances in at least one sample) presumed to include indigo-reducing bacteria. All data represent the average values from two independent cultures and error bars represent standard deviations.

The PCA based on microbial community profiles clearly demonstrated that the microbial communities in the dye suspensions were altered by the addition of commercial *sukumo* (Figs. 4C and 4F). The relative abundances of several phylotypes varied markedly depending on the type of immature *sukumo* used (T-12w or W-8w) and the presence or absence of commercial *sukumo* supplementation (Fig. S1). Most of them were phylotypes presumed to include indigo-reducing bacteria, which are shown in Fig. 4G. Among them, g_*Corynebacterium* was predominantly observed in T-12w cultures, particularly during the later stages, whereas it constituted only a minor fraction in W-8w cultures. This observation is consistent with the community analysis of the *sukumo* fermentation process, in which g_*Corynebacterium* was predominantly present only during Phases 3–4 of T *sukumo* fermentation (Fig. 3A). In contrast, the relative abundances of g_*Pseudomonas*, g_*Oceanobacillus*, and g_*Amphibacillus* were higher in W-8w cultures than in T-12w cultures. Notably, g_*Oceanobacillus* showed a higher relative abundance in W-8w+*sukumo* than in W-8w, particularly during the initial phase (weeks 1 to 2), when an improvement in dyeing performance was observed. g_*Alkalibacillus* was dominant in both T-12w and W-8w cultures, but its abundance was lower in the respective +*sukumo* cultures. The population dynamics of f_*Tissierellaceae* exhibited intriguing behavior: it was detected only in the cultures supplemented with commercial *sukumo* within the T-12w culture series. In contrast, within the W-8w culture series, its abundance was lower in the cultures supplemented with commercial *sukumo*. These observed shifts in the population dynamics of indigo-reducing bacteria are presumed to have contributed to the enhanced dyeing performance of dye suspensions by the addition of commercial *sukumo*.

It should be noted that when more immature *sukumo* with a shorter fermentation duration, specifically, week 8 of T *sukumo* and week 4 of W *sukumo*, was used, the dye suspensions exhibited almost no staining activity, regardless of the addition of commercial *sukumo* (data not shown). Even at this early stage of fermentation, the immature *sukumo* is expected to contain a sufficient amount of indigo (Fig. 2B), and the addition of commercial *sukumo* should provide an adequate microbial source. These observations suggest that factors other than indigo content and its function as a microbial inoculum may also contribute to determining dyeing performance.

### *Effect of supplementation of an indigo-reducing bacterium on dye suspensions prepared with immature* sukumo

In the experiment shown in Fig. 4, two potential confounding factors must be considered: substantial shifts in the microbial community structure and the possible influence of indigo and organic compounds introduced by the commercial *sukumo*. To evaluate the direct effect of supplementation of indigo-reducing bacteria, we conducted an additional experiment in which dye suspensions were prepared using immature *sukumo* and supplemented with a pure culture of the indigo-reducing bacterium, namely *Tissierellaceae* strain TU-1. This strain exhibits exceptionally high indigo-reducing activity in pure culture (Tu *et al*., 2025) and *Tissierellaceae* bacteria were presumed to have contributed to the improved dyeing performance observed in the T-12w+*sukumo* cultures as shown in Fig. 4. It should be noted that the immature *sukumo* used in this experiment (T’-12w) was produced in a different year from that used in the previous experiments.

The addition of a culture of *Tissierellaceae* strain TU-1 to dye suspensions prepared with immature *sukumo* resulted in a trend of improved dyeing performance (Fig. 5A). Quantitative analysis of staining intensity revealed significant increases during the early (weeks 1 and 2) and late (weeks 5 and 6) stages of cultures (Fig. 5B). The PCA based on microbial community profiles indicated that the addition of *Tissierellaceae* strain TU-1 had minimal impact on the overall microbial structure (Fig. 5C). This was further supported by the population dynamics of dominant phylotypes (Fig. S2), which showed that, unlike the case with commercial *sukumo* supplementation, the abundances of dominant phylotypes remained largely unchanged. Notably, no significant increases or decreases in known indigo-reducing bacteria other than *Tissierellaceae* were observed by adding the culture of strain TU-1.

**Fig. 5.**
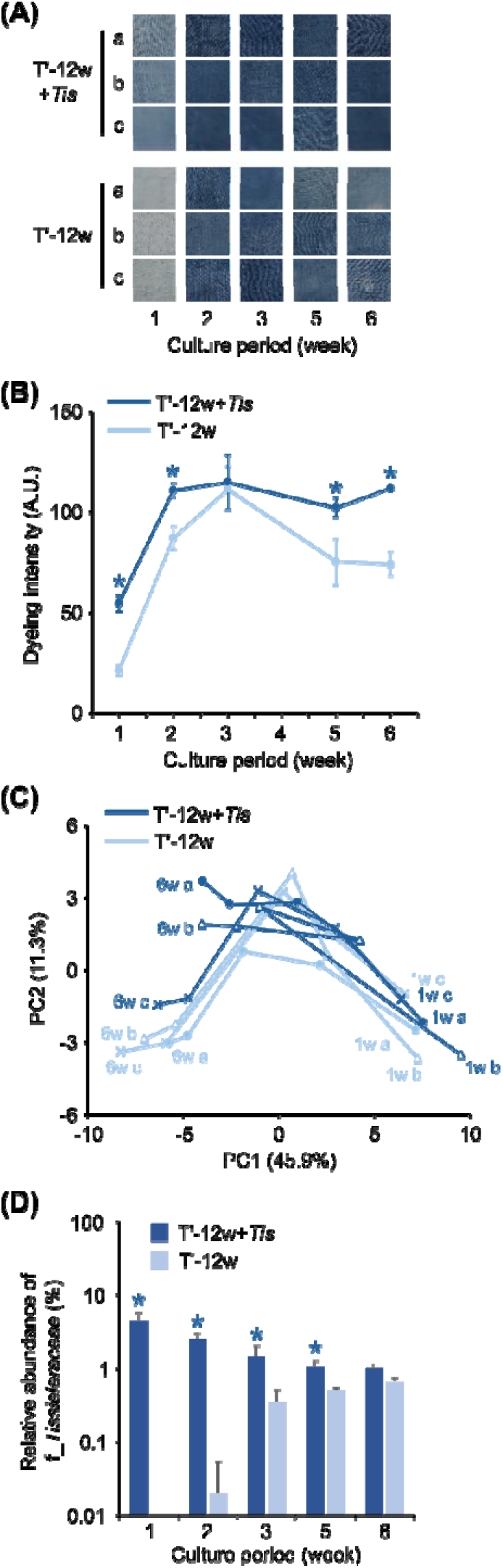
Culture experiments of dye suspensions using immature *sukumo* to evaluate the effect of supplementation with an indigo-reducing bacterium. The immature *sukumo* used for the T’-12w culture was collected at week 12 of T *sukumo* production. T’-12w+*Tis* indicates cultures supplemented with a pure culture of the indigo-reducing bacterium *Tissierellaceae* strain TU-1. (A) Photographs of cotton fabrics dyed using each dye suspension culture. Labels ‘a’ to ‘c’ represent three independent culture replicates. (B) Quantification of color intensity based on the dyed fabrics shown in (A). Asterisks indicate significantly higher color intensity in the +*Tis* cultures, which was assessed by Student’s *t*-test. (C) Principal component analysis based on microbial community profiles of the dye suspension cultures, as presented in Supplementary Fig. S2. Data points corresponding to weeks 1 and 6 of each culture are annotated. (G) Temporal changes in the relative abundance of f_*Tissierellaceae*. Asterisks indicate significantly higher abundance in the +*Tis* cultures, which was assessed by Student’s *t*-test. All data represent the mean values from three independent cultures, and error bars indicate standard deviations.

The relative abundance of *Tissierellaceae* (Fig. 5D; see also f_*Tissierellaceae* in Fig. S2) differed markedly during the early culture period (weeks 1 and 2). Given the minimal changes in the relative abundances of other microbial taxa, the observed improvement in staining intensity during the early phase is likely attributable to the activity of *Tissierellaceae* strain TU-1. In contrast, enhanced staining was also observed during the later culture phase (weeks 5 and 6), when the difference in the relative abundance of *Tissierellaceae* between the two conditions was no longer pronounced. This result suggests that staining activity cannot be explained solely by the quantity of indigo-reducing bacteria. For example, it has been reported that the activity of indigo-reducing bacteria, including strain TU-1, strongly depends on the production of electron mediators, such as insoluble quinone compounds, that facilitate electron transfer to insoluble indigo (Nakagawa *et al*., 2024; Tu and Yumoto, 2025). The improved staining activity observed in this experiment may therefore be attributed to factors other than microbial abundance, such as the concentration of electron mediators present in the dye suspension.

## Conclusion

In this study, we conducted a time-series analysis of the microbial communities during the *sukumo* production process, along with laboratory culture experiments of dye suspension using immature *sukumo*. Our findings revealed that *sukumo* plays a critical role not only as a pigment source but also as a microbial source in traditional indigo dyeing. In addition, although present at low relative abundances, multiple indigo-reducing bacteria began to emerge during the later stages of *sukumo* production, suggesting that the extended four-month production period is essential for nurturing these functional microbes. Culture experiments using immature *sukumo* further indicated that modification of the microbial community, such as through the addition of beneficial strains, may improve staining activity of dye suspensions. Further microbiological research will deepen our understanding of this traditional fermentation-based dyeing process and highlight the rare and compelling example of microorganisms playing an active role in the world of clothing and fashion.

## Data availability

The sequence data obtained in the present study have been deposited in DDBJ/EMBL/GenBank under the accession numbers xxxxxx.

## Acknowledgement

The authors express their appreciation to Mr. Osamu Nii, an Aizome master of Nii Seiaisyo for his invaluable suggestions and instruction regarding Japanese traditional dyeing techniques and *sukumo* production. The authors would like to express appreciation to Dr. Zhihao Tu, Ms. Ai Miura, Ms. Mika Yamamoto, and Ms. Nana Miyazaki for technical assistance.

## References

Agafonova, N.V., Belova, A.A., Kaparullina, E.N., Tarlachkov, S.V., Kopitsyn, D.S., Machulin, A.V., and Doronina, N.V. (2023) *Ancylobacter radicis* sp. nov., a novel aerobic methylotrophic bacteria associated with plants. Antonie Van Leeuwenhoek 116: 855–866.

Aino, K., Narihiro, T., Minamida, K., Kamagata, Y., Yoshimune, K., and Yumoto, I. (2010) Bacterial community characterization and dynamics of indigo fermentation. FEMS Microbiol Ecol 74: 174–183.

Aino, K., Hirota, K., Okamoto, T., Tu, Z., Matsuyama, H., and Yumoto, I. (2018) Microbial communities associated with indigo fermentation that thrive in anaerobic alkaline environments. Front Microbiol 9: 2196.

Bolyen, E., Rideout, J.R., Dillon, M.R., Bokulich, N.A., Abnet, C.C., Al-Ghalith, G.A., et al. (2019) Reproducible, interactive, scalable and extensible Microbiome data science using QIIME 2. Nat. Biotechnol. 37: 852–857.

de Maayer, P., Chan, W.Y., Rubagotti, E., Venter, S.N., Toth, I.K., Birch, P.R.J., and Coutinho, T.A. (2014) Analysis of the *Pantoea ananatis* pan-genome reveals factors underlying its ability to colonize and interact with plant, insect and vertebrate hosts. BMC Genom 15: 1–28.

Dobrzyński, J., Wróbel, B., and Górska, E.B. (2023) Taxonomy, ecology, and cellulolytic properties of the genus *Bacillus* and related genera. Agriculture 13: 1979.

Farjana, N., Tu, Z., Furukawa, H., and Yumoto, I. (2023) Environmental factors contributing to the convergence of bacterial community structure during indigo reduction. Front Microbiol 14: 1097595.

Han, S.I., Lee, J.C., Lee, H.J., and Whang, K.S. (2013) *Planifilum composti* sp. nov., a thermophile isolated from compost. Int J Syst Evol Microbiol 63: 4557–4561.

Haruta, S., Nakayama, T., Nakamura, K., Hemmi, H., Ishii, M., Igarashi, Y., and Nishino, T. (2005) Microbial diversity in biodegradation and reutilization processes of garbage. J Biosci Bioeng 99: 1–11.

Hayakawa, M., Yamamura, H., Nakagawa, Y., Kawa, Y., Hayashi, Y., Misonou, T., et al. (2010) Taxonomic diversity of Actinomycetes isolated from swine manure compost. Actinomycetologica 24: 58–62.

Hirota, K., Aino, K., Nodasaka, Y., Morita, N., and Yumoto, I. (2013a) *Amphibacillus indicireducens* sp. nov., an alkaliphile that reduces an indigo dye. Int J Syst Evol Microbiol 63: 464–469.

Hirota, K., Aino, K., Nodasaka, Y., and Yumoto, I. (2013b) *Oceanobacillus indicireducens* sp. nov., a facultative alkaliphile that reduces an indigo dye. Int J Syst Evol Microbiol 63: 1437–1442.

Hirota, K., Aino, K., and Yumoto, I. (2013c) *Amphibacillus iburiensis* sp. nov., an alkaliphile that reduces an indigo dye. Int J Syst Evol Microbiol 63: 4303–4308.

Hirota, K., Hanaoka, Y., Nodasaka, Y., and Yumoto, I. (2013d) *Oceanobacillus polygoni* sp. nov., a facultatively alkaliphile isolated from indigo fermentation fluid. Int J Syst Evol Microbiol 63: 3307–3312.

Hugerth, L.W., Wefer, H.A., Lundin, S., Jakobsson, H.E., Lindberg, M., Rodin, S., Engstrand, L., and Andersson, A.F. (2014) DegePrime, a program for degenerate primer design for broad-taxonomic-range PCR in microbial ecology studies. Appl Environ Microbiol 80: 5116–5123.

Iguchi, H., Yurimoto, H., and Sakai, Y. (2015) Interactions of methylotrophs with plants and other heterotrophic bacteria. Microorganisms 3: 137–151.

Janda, J.M., and Abbott, S.L. (2021) The changing face of the family *Enterobacteriaceae* (order: “*Enterobacterales*”): new members, taxonomic issues, geographic expansion, and new diseases and disease syndromes. Clin Microbiol Rev 34: e00174–20.

Li, S., Shi, Y., Huang, H., Tong, Y., Wu, S., and Wang, Y. (2022) Fermentation blues: analyzing the microbiota of traditional indigo vat dyeing in Hunan, China. Microbiol Spectr 10: e0166322.

Liu, Q., He, X., Luo, G., Wang, K., and Li, D. (2022) Deciphering the dominant components and functions of bacterial communities for lignocellulose degradation at the composting thermophilic phase. Bioresour Technol 348: 126808.

Lopes, H.F.S., Tu, Z., Sumi, H., and Yumoto, I. (2021) Analysis of bacterial flora of indigo fermentation fluids utilizing composted indigo leaves (*sukumo*) and indigo extracted from plants (Ryukyu-ai and Indian indigo). J Biosci Bioeng 132: 279–286.

Nakagawa, K., Takeuchi, M., Tada, M., Matsunaga, M., Kugo, M., Kiyofuji, S., et al. (2022) Isolation and characterization of indigo-reducing bacteria and analysis of microbiota from indigo fermentation suspensions. Biosci Biotechnol Biochem 86: 273–281.

Nakagawa, K., Ohata, H., Takeuchi, M., Matsunaga, M., Sowa, K., Sakamoto, T., et al. (2024) Effects of lignin on indigo-reducing activity and indigo particle size in indigo dye suspensions. Biosci Biotechnol Biochem 89: 141–144.

Nicholson, S.K., and John, P. (2005) The mechanism of bacterial indigo reduction. Appl Microbiol Biotechnol 68: 117–123.

Nishita, M., Hirota, K., Matsuyama, H., and Yumoto, I. (2017) Development of media to accelerate the isolation of indigo-reducing bacteria, which are difficult to isolate using conventional media. World J Microbiol Biotechnol 33: 133.

Park, S., Ryu, J.Y., Seo, J., and Hur, H.G. (2012) Isolation and characterization of alkaliphilic and thermotolerant bacteria that reduce insoluble indigo to soluble leuco-indigo from indigo dye vat. J Korean Soc Appl Biol Chem 55: 83−88.

Riahi, H.S., Heidarieh, P., and Fatahi-Bafghi, M. (2022) Genus *Pseudonocardia*: What we know about its biological properties, abilities and current application in biotechnology. J Appl Microbiol 132: 890–906.

Tu, Z., Lopes, H.F.S., Igarashi, K., and Yumoto, I. (2019) Characterization of the microbiota in long- and short-term natural indigo fermentation. J Ind Microbiol Biotechnol 46: 1657–1667.

Tu, Z., Lopes, H.F.S., Narihiro, T., and Yumoto, I. (2021) The mechanism underlying of long-term stable indigo reduction state in indigo fermentation using *Sukumo* (composted *Polygonum tinctorium* leaves). Front Microbiol 12: 698674.

Tu, Z., and Yumoto, I. (2025) Isolation of a *Tissierellaceae* bacterium exhibiting a high reduction potential for insoluble indigo dyes. Microbes Environ 40: ME24104.

Ueno, Y. (2023) Indigo dyeing and fermentation sukumo, essential for traditional Japanese aizome. Art Sci 7: 1–7.

Watanabe, K., Nagao, N., Toda, T., and Kurosawa, N. (2008) Changes in bacterial communities accompanied by aggregation in a fed-batch composting reactor. Curr Microbiol 56: 458–467.

Watanabe, M., Kojima, H., and Fukui, M. (2015) *Limnochorda pilosa* gen. nov., sp. nov., a moderately thermophilic, facultatively anaerobic, pleomorphic bacterium and proposal of *Limnochordaceae* fam. nov., *Limnochordales* ord. nov. and *Limnochordia* classis nov. in the phylum Firmicutes. Int J Syst Evol Microbiol 65: 2378–2384.

Wei, H., Wang, L., Hassan, M., and Xie, B. (2018) Succession of the functional microbial communities and the metabolic functions in maize straw composting process. Bioresour Technol 256: 333–341

Yang, G., and Zhou, S. (2014) *Sinibacillus soli* gen. nov., sp. nov., a moderately thermotolerant member of the family *Bacillaceae*. Int J Syst Evol Microbiol 64: 1647–1653.

Zhou, H., Di, L., Hua, X., Deng, T., and Wang, X. (2023) Bacterial community drives the carbon source degradation during the composting of *Cinnamomum camphora* leaf industrial extracted residues. Microbiol Res 14: 229–242.

