## Supplementary Fig S1, S2 for "The significance of *sukumo* (composted indigo leaves) as a microbial source for traditional Japanese indigo dyeing"

**Supplementary materials**

Figs. S1 and S2

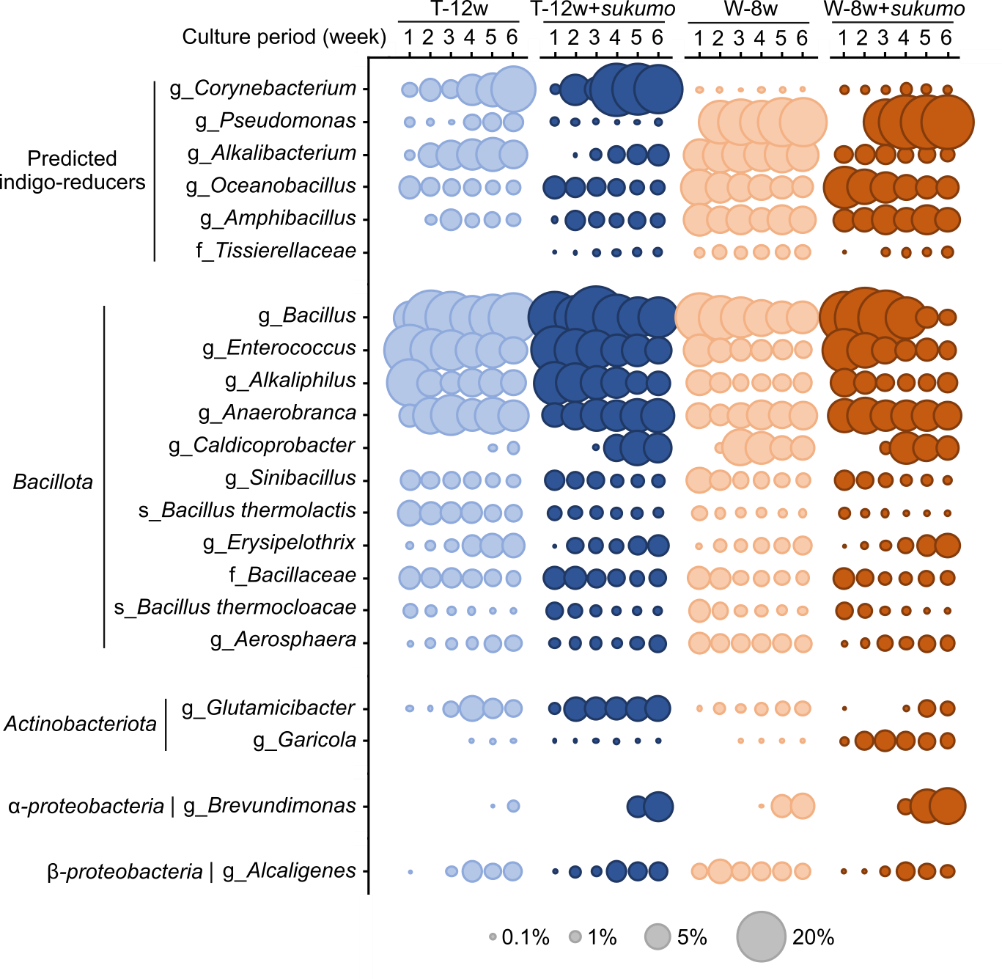
**Fig. S1.** Microbial community dynamics of dye suspension cultures using immature *sukumo* supplemented with commercial *sukumo* as a microbial source. Immature *sukumo* samples were collected from the production process immediately after the pH reached 8. T-12w and W-8w refer to dye suspension cultures using 12-week of T *sukumo* and 8-week of W *sukumo*, respectively. T-12w+*sukumo* and W-8w+*sukumo* indicate cultures supplemented with a 1/20 volume of commercially available mature *sukumo* as a microbial source. Temporal changes in the relative abundance of dominant phylotypes (defined as those comprising >3% in at least one sample) are presented.

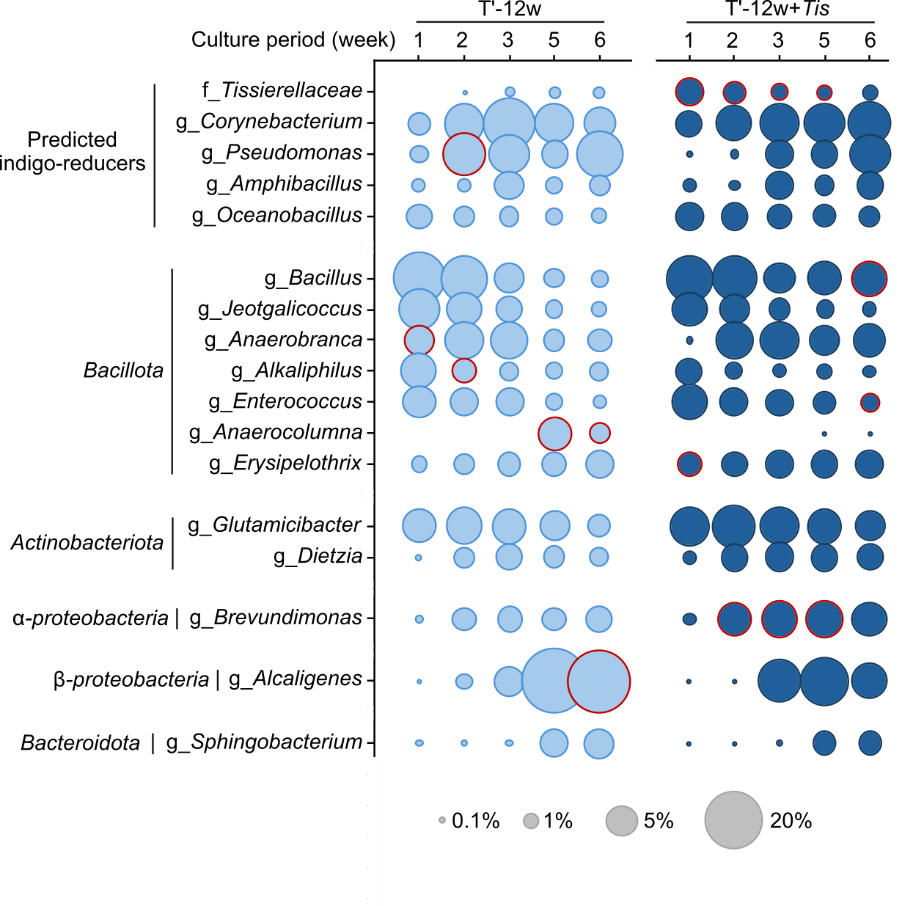
**Fig. S2.** Microbial community dynamics of dye suspension cultures using immature *sukumo* supplemented with an indigo-reducing bacterium. The immature *sukumo* used for the T’-12w culture was collected at week 12 of T *sukumo* production. T’-12w+*Tis* indicates cultures supplemented with a pure culture of the indigo-reducing bacterium *Tissierellaceae* strain TU-1. Temporal changes in the relative abundance of dominant phylotypes (defined as those comprising >3% in at least one sample) are presented. Red-outlined circles indicate phylotypes with significantly higher relative abundance under the corresponding conditions (>2-fold change, *p* < 0.05, and >1% relative abundance).
